# Pan-cancer metabolic landscapes: A multi-omics view

**DOI:** 10.1101/2025.08.14.670283

**Authors:** Yiguang Bai, Qiaoling Chen, Yuan Li

## Abstract

Metabolic reprogramming fuels cancer progression, but whether common metabolic patterns exist across tumor types remains elusive. To address this, we developed parseMetab, an R package that integrates large-scale proteomic, transcriptomic, and spatial transcriptomic datasets from 24 human cancers and 3,226 samples to map pan-cancer metabolic dysregulation. Strikingly, glycan biosynthesis pathways – especially those driving fucosylation and sialylation – were consistently upregulated, while early sugar nucleotide precursors were suppressed. Nucleotide metabolism was broadly enhanced, uncovering conserved metabolic programs that enable tumor growth, adaptation, and immune evasion. Our results highlight universal metabolic vulnerabilities that may be therapeutically exploited across diverse cancer contexts.

**Significance statement:** While metabolic reprogramming is a known hallmark of cancer, identifying universal patterns across diverse tumor types remains a major challenge. We developed **parseMetab**, a computational framework to integrate large-scale proteomic and transcriptomic data from over 3,200 samples across 24 human cancers. Our analysis reveals a striking, conserved “glyco-switch”: tumors consistently prioritize the production of complex, immune-evading surface sugars (fucosylation and sialylation) while depleting their own internal sugar precursor pools. Furthermore, we demonstrate that nucleotide metabolism is broadly hyperactivated across nearly all studied malignancies. By defining these ubiquitous metabolic signatures, this study uncovers conserved vulnerabilities that transcend specific cancer types, providing a map for developing broad-spectrum metabolic therapies to combat tumor growth and immune evasion.

## Introduction

Metabolic reprogramming is a hallmark of cancer, enabling malignant cells to proliferate rapidly, survive stress, and adapt to dynamic microenvironments [1–3]. Beyond altered energy production, cancer metabolism supports biosynthesis, redox balance, and signaling pathways that collectively drive tumor growth [4, 5]. While central metabolic pathways have been extensively studied, key processes such as glycan biosynthesis – which play critical roles in protein modification, cell communication, and immune interactions – have received comparatively limited pan-cancer attention [6]. This is notable given that glycan synthesis – a complex enzymatic process occurring within the endoplasmic reticulum and Golgi apparatus – adds regulatory layers that significantly influence tumor behavior [7, 8]. Although aberrant glycosylation is broadly recognized as a hallmark of tumorigenesis affecting essential cancer cell functions, its prevalence and variation across diverse tumor types remain incompletely characterized in systematic pan-cancer studies [9].

Many studies of cancer metabolism focus on individual tumor types [1–3]. Pan-cancer analyses, when performed, are frequently based on a single omics layer – most commonly transcriptomic data from The Cancer Genome Atlas (TCGA) [10–13]. Although multi-omics pan-cancer approaches have emerged, they often emphasize selected metabolic pathways rather than systematically characterizing the full metabolic landscape [14–17]. Here, we uniquely integrate independent proteomic, transcriptomic, and spatial transcriptomic dataset collections to construct a multi-dimensional pan-cancer metabolic landscape. By applying complementary computational approaches across diverse omics modalities and leveraging spatial data to distinguish tumor microenvironment compartments, we offer a more comprehensive and biologically contextualized view of tumor metabolism. This integrated framework facilitates the identification of shared metabolic programs and highlights pathways that may serve as broadly targetable vulnerabilities across cancer types.

## Results and Discussion

To characterize pan-cancer metabolic reprogramming, tumor and matched non-tumor samples were analyzed across independent proteomics, bulk transcriptomics, and spatial transcriptomics datasets, encompassing 24 cancer types and 3,226 samples (1,610 tumor, 1,616 non-tumor) from multiple studies (Fig. 1A, Table S1). Metabolic task activities were first inferred using the CellFie framework on two proteomics datasets: the unified pan-cancer resource by Zhou et al. [18] and a curated proteomics collection by Hu et al. [19], identifying 176 and 138 active tasks, respectively, grouped into seven metabolic classes comprising 77-136 proteins (Fig. 1B; Table S1). Complementarily, Gene Set Variation Analysis (GSVA) [20] was performed using the Kyoto Encyclopedia of Genes and Genomes (KEGG) database [21]. GSVA was applied to the aforementioned proteomics datasets, transcriptomic data from The Cancer Genome Atlas (TCGA) [22], and spatial transcriptomic pseudobulk RNA-seq data from SMTdb [23]. This analysis resolved 55–83 metabolic pathways across eleven classes, representing 1,454–1,705 genes (Fig. 1B, S1Q-S; Tables S1, S2).

**Fig. 1.**
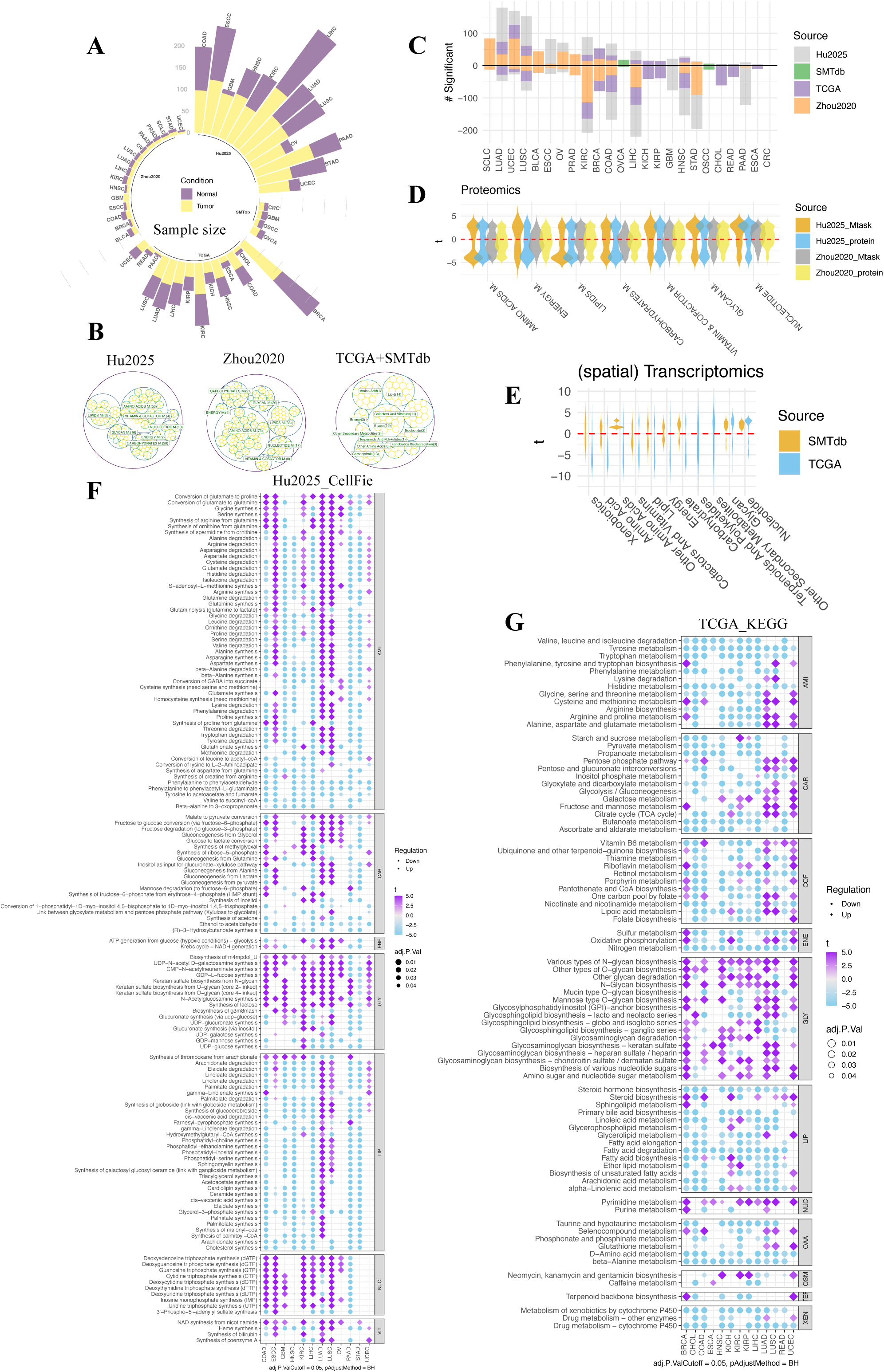
Pan-cancer metabolic dysregulation across independent pan-cancer datasets from distinct omics modalities. **(A)** Sample sizes for the four independent pan-cancer datasets analyzed. **(B)** Overview of the number of identified metabolic tasks/pathways per metabolic class, in each dataset. **(C)** Number of significantly up- or downregulated tasks/pathways per dataset: Hu2025 (adjusted P-value < 0.05), Zhou2020 (adj. P < 0.1), TCGA (adj. P < 0.05), and SMTdb (P < 0.1). Downregulated features are plotted as negative values. **(D/E)** Violin plots showing the distribution of moderated *t* statistics from task/pathway- or protein-level differential analyses for each metabolic class using either the proteomics (D) or the bulk/spatial transcriptomics (E) datasets. **(F/G)** Dot plots of task/ pathway -level dysregulation across cancer types in either Hu2025 proteomics (F) or TCGA transcriptomics (G) data. Here in this figure, the metabolic tasks are CellFie-based for the two proteomics datasets (Hu2025 & Zhou2020), the metabolic pathways are KEGG-based for the TCGA and SMTdb datasets. In (F) and (G), color represents moderated *t* statistics, and dot size reflects adjusted *p* values; only significant features (adj. P < 0.05) are shown.

Comparison of tumors with matched normal samples revealed widespread metabolic reprogramming across cancer types, with consistent patterns across independent datasets. Notably, Uterine Corpus Endometrial Carcinoma (UCEC) showed strong metabolic upregulation, potentially driven by frequent activation of the insulin-phosphoinositide 3 kinase (PI3K) signaling pathway and by estrogen signaling [24] (Fig. 1C, 1F, 1G and S2A). In contrast, Liver Hepatocellular Carcinoma (LIHC) consistently ranked among the most metabolically downregulated cancers (Fig. 1C, 1F, 1G and S2A), in line with a liver tissue metabolomics study showing that the majority of significantly altered metabolites were reduced in tumor tissue relative to matched adjacent liver [25]. These results highlight robust and reproducible pan-cancer metabolic alterations across diverse cohorts.

Metabolic reprogramming in cancer showed significant heterogeneity across metabolic classes. Notably, nucleotide and glycan metabolism were consistently upregulated across all four datasets, with stronger effects observed in (spatial) transcriptomics (Fig. 1D-E, Fig. S1T-U, Table S3). This pattern was further confirmed at the protein level in the two proteomics datasets using paired differential analysis (Fig. 1D, Table S3). Multi-omics integration using iNMF – restricted to glycan and nucleotide pathways – demonstrated a clear separation between malignant and normal tissues, while successfully mitigating batch effects across the integrated omics layers (Fig. 2A-B). Conversely, amino acid metabolism was among the most downregulated classes in three bulk omics comparisons (tumor vs. matched normal), a trend not seen in spatial omics comparing malignant and stromal spots (Fig. 1D-E, Fig. S1T-U).

**Fig. 2.**
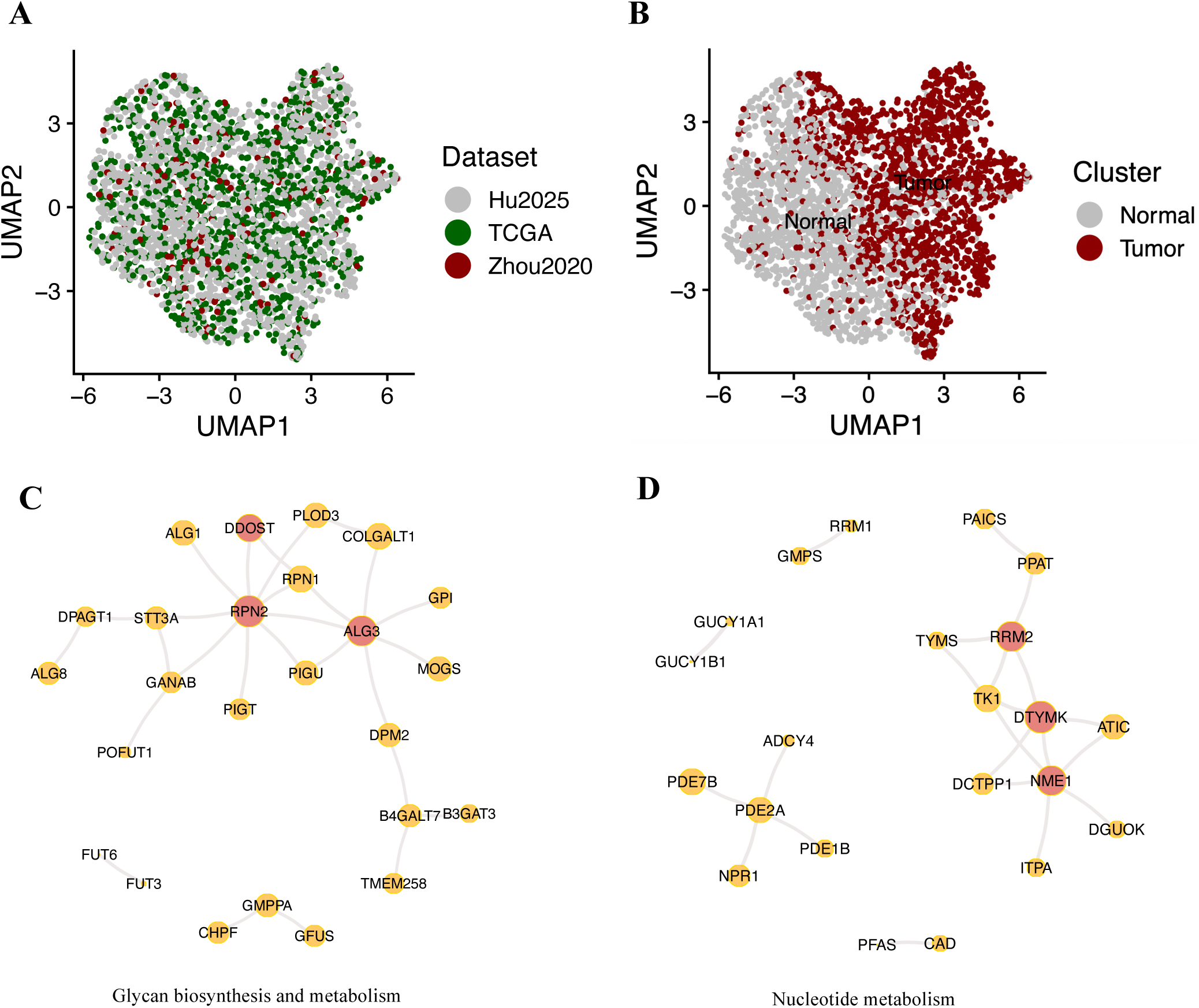
Multi-omics integration and gene co-expression networks for glycan and nucleotide metabolism. (A-B) Integrative visualization of bulk proteomics (Hu2025 & Zhou2020) and bulk transcriptomics (TCGA) datasets. (C-D) Gene co-expression networks for glycan and nucleotide metabolism. Nodes represent genes, sized by normalized connectivity, with the top 1% most connected genes (≥3 per class) highlighted as hubs (red). Edges show absolute correlation > 0.6, width scaled by correlation strength; grey edges indicate positive correlations and blue edges indicate negative correlations. Networks were visualized using a Fruchterman-Reingold layout.

### The tumor glyco-switch: from simple sugars to complex capping

Among those broadly conserved alterations, glycan metabolism displayed a particularly coherent and biologically informative pattern. Tumors showed coordinated upregulation of pathways generating activated sugars for complex glycan modifications – including fucosylation and sialylation – accompanied by downregulation of common sugar nucleotide precursors such as uridine diphosphate glucose (UDP-glucose) and guanosine diphosphate mannose (GDP-mannose) (Fig. 1F-G, S2A-B, S3A-B, Table S3). Consistently, perturbations in mannose metabolism have been shown to disrupt nucleotide sugar biosynthesis and impair cancer cell proliferation [26, 27]. This shift is consistent with cancer-associated remodeling of cell surface, secreted, and extracellular matrix glycoproteins, which enhances invasiveness, immune evasion, and signaling through structurally complex glycans [8, 28]. The reduced synthesis of early sugar precursors likely reflects a metabolic prioritization toward downstream glycosylation pathways that promote tumor survival and progression.

Biologically, increased CMP-N-acetylneuraminate and guanosine diphosphate L-fucose (GDP-L-fucose) synthesis facilitate the generation of sialylated and fucosylated glycans that modulate immune recognition and metastasis [29, 30]. Concurrently, the reduction in UDP-glucose and GDP-mannose production may indicate altered N-glycan core assembly or glycogen metabolism, further supporting a tumor-adapted metabolic state.

This robust, cross-cancer evidence – derived from a unified analytical framework – strengthens the concept that alterations in glycosylation pathways represent a broadly conserved metabolic hallmark of tumorigenesis. While previous studies have suggested glycosylation as a universal feature of malignant cells [31], our systematic analysis provides the first comprehensive, multi-dimensional confirmation of this phenomenon across diverse cancers, underscoring its potential as a target for diagnostic and therapeutic strategies [32]. Gene co-expression network analysis in glycan metabolism identified key hub genes – e.g. ALG3 (Alpha-1,3-mannosyltransferase), RPN2 (ribophorin II), and DDOST (Dolichyl-diphosphooligosaccharide – protein glycosyltransferase 48 kDa subunit) – that may drive this metabolic rewiring and represent candidates for targeted cancer therapies (Fig. 2C-D). Notably, ALG3, an ER-localized glycosylation enzyme, is implicated in promoting tumor growth, invasion, and resistance to radio-, and immunotherapies [33–35]. Moreover, RPN2, an oncogenic subunit of the N-oligosaccharyltransferase complex, is aberrantly expressed in multiple human cancers, where it promotes tumor progression, metastasis, and drug resistance [36].

### Pan-cancer upregulation of nucleotide metabolism: a systemic hallmark

Moreover, nucleotide metabolism – covering the synthesis, salvage, and catabolism of purine and pyrimidine nucleotides – is a fundamental metabolic vulnerability exploited by cancer cells to drive proliferation, evade immune surveillance, resist chemotherapy, and promote metastasis [37, 38]. Leveraging a comprehensive pan-cancer analytical framework, we systematically quantify and characterize the upregulation of nucleotide metabolic pathways across a wide spectrum of tumor types. Our findings reveal that this metabolic hallmark is not confined to isolated cancer subsets but represents a widespread feature observed in multiple malignancies, thereby expanding upon and unifying prior reports focused on individual tumor contexts [39].

### Beyond the hallmark: the heterogeneous metabolic architecture of cancer

In contrast to the consistent upregulation observed in nucleotide and glycan metabolism, our analysis reveals that the dysregulation of other metabolic pathways exhibits substantial heterogeneity both between different cancer types and across independent dataset collections. This variability is likely driven by a complex interplay of factors, including tissue-specific metabolic programming, diverse oncogenic mutations, the influence of the tumor microenvironment, and technical differences in sample processing and data generation [3, 40]. Tissue origin profoundly shapes the metabolic phenotype of tumors, as distinct organs possess unique baseline metabolic demands and enzyme expression profiles that tumors may retain or reprogram selectively [41]. Furthermore, oncogenic drivers such as MYC, KRAS (Kristen Rat Sarcoma viral oncogene), and TP53 (tumor protein p53) mutations modulate metabolic pathways differently across cancer types, resulting in context-dependent metabolic adaptations [2]. The tumor microenvironment, including nutrient availability, hypoxia, and stromal interactions, further sculpts metabolic reprogramming in a spatially and temporally dynamic manner [42]. Technical heterogeneity across datasets – including differences in sample collection, sequencing platforms, and bioinformatic pipelines – can also contribute to observed variability, underscoring the need for careful data integration and validation [43]. This pronounced metabolic heterogeneity challenges the identification of universal metabolic vulnerabilities beyond the consistently conserved nucleotide and glycan pathways, emphasizing the value of integrative multi-cohort analyses to comprehensively capture metabolic diversity and enhance the robustness and predictive power of metabolic biomarkers.

## Conclusion

Our integrative analysis across proteomics, bulk transcriptomics, and spatial transcriptomics reveals conserved pan-cancer metabolic reprogramming, with coordinated upregulation of nucleotide and glycan metabolism and key hubs such as ALG3 and RPN2. These results highlight broadly conserved metabolic vulnerabilities and the importance of glycosylation in tumor progression. Nevertheless, limitations including tumor purity and the observational nature of multi-omics data underscore the need for future validation using direct metabolomics/glycomics measurements and functional experiments.

## Methods

To study pan-cancer metabolic reprogramming, we compared tumor and matched non-tumor samples across multi-omics datasets, integrating two proteomics resources (Hu2025, Zhou2020) [18–19] alongside the TCGA transcriptomics cohort [22] and SMTdb spatial transcriptomics collection [23]. For bulk datasets, tumor samples were compared to matched non-tumor tissues, while for spatial transcriptomics data, comparisons were conducted between malignant spots and stromal spots within the same tissue section. For each STMdb tissue section, raw RNA counts from all malignant and stromal spots were aggregated separately to generate pseudobulk RNA-seq profiles. The majority of analyzed samples were matched tumor-normal pairs from the same patients, enabling robust intra-patient comparisons across diverse cancer contexts (Table S1).

We used the Zhou et al. dataset in its preprocessed form as provided by the original publication [18], while the Hu et al. collection was curated and uniformly reprocessed by the authors [19]. Transcriptomic data for multiple cancer types were obtained from TCGA [22] using TCGAbiolinks v2.34.1. Raw gene-level counts generated using Spliced Transcripts Alignment to a Reference (STAR) were retrieved, including only Primary Tumor and Solid Tissue Normal samples with paired data per subject. Genes were filtered to retain those with counts > 5 in at least 15% of samples per group or at least two samples when 15% was < 2. Data were normalized using the limma-voom pipeline (limma v3.62.1[44], edgeR v4.4.0 [45]), yielding log2-transformed counts per million (logCPM). Spatial transcriptomics data for multiple cancer types were obtained from SMTdb [23]. For each cancer type, expression data per spot and spatial annotations (malignant, stromal, boundary) were retrieved from the SMTdb website. We included only cancer types with at least three slices containing malignant spots and three with stromal spots. For each slice, raw counts from malignant and stromal spots were aggregated separately into pseudobulk profiles, typically yielding one malignant and one stromal sample per slice. Genes were filtered using the same criteria as TCGA data. Pseudobulk data were normalized with edgeR (v4.4.0) using TMMwsp normalization, which accounts for sample composition bias and variable expression across distinct cell populations, resulting in logCPM values.

### Metabolic task scoring using CellFie

To interpret proteomics data across 16 cancer types from the Hu2025 and Zhou2020 datasets, proteins were grouped into metabolic tasks, and task activities were quantified using CellFie on the GenePattern webserver [46]. Metabolic tasks represent small reaction modules converting specific substrates to products, reflecting metabolic reprogramming that supports tumor growth and therapy resistance [2,3].

Proteins were mapped to Entrez Gene IDs using org.Hs.eg.db v3.20.0 and AnnotationDbi v1.68.0. The Zhou2020 dataset was log[J-transformed prior to analysis, while the Hu2025 dataset was exponentiated to avoid negative values. Feature activity was determined via local thresholding (Local.Threshold.Type = “minmaxmean”), with thresholds set as feature means constrained between the 25th and 75th percentiles of overall abundance.

All Zhou2020 samples were analyzed together, whereas Hu2025 datasets were submitted individually. CellFie categorized metabolic tasks into seven classes (lipid, energy, nucleotide, amino acid, glycan, carbohydrate, and vitamin & cofactor metabolism). To align with Kyoto Encyclopedia of Genes and Genomes (KEGG) classifications, several tasks originally assigned to carbohydrate metabolism were reassigned to glycan metabolism: Synthesis of lactose, UDP-glucose synthesis, UDP-galactose synthesis, UDP-glucuronate synthesis, GDP-L-fucose synthesis, GDP-mannose synthesis, UDP-N-acetyl D-galactosamine synthesis, CMP-N-acetylneuraminate synthesis, N-Acetylglucosamine synthesis, Glucuronate synthesis (via inositol), Glucuronate synthesis (via UDP-glucose).

### Metabolic enrichment scoring using KEGG-based GSVA

To infer metabolic activity from the two aforementioned proteomics, as well as the TCGA transcriptomic and SMTdb pseudobulk RNA-seq data, metabolic gene sets/pathways and their classes were extracted from KEGG using KEGGREST v1.46.0 [47]. Gene Set Variation Analysis (GSVA) was performed on the metabolic gene sets (size 5–500) to calculate pathway-level enrichment scores reflecting relative metabolic activity per sample. GSVA, a non-parametric unsupervised method, estimates variation in pathway activity across samples by integrating gene expression into pathway-centric scores, enabling sample-wise comparisons. Analysis was conducted using GSVA v2.0.7 [20].

### Differential analysis of metabolic activities between cancer and normal samples

To understand how cancer cells reprogram their metabolic processes to support rapid growth, survival, invasion, and therapy resistance, we performed univariate differential analysis using linear models implemented in limma v3.62.2. Each metabolic task/pathway was tested individually for differences between tumor and matched normal samples. In this analysis, each cancer type was analyzed separately using the Zhou2020, Hu2025, Zhou2020_KEGG, Hu2025_KEGG, TCGA, and SMTdb dataset collections.

Differential analysis was only performed on cancer types with both cancer and normal samples, and there were at least two Cancer-Normal pairs, paired differential analysis was conducted using limma v.3.62.1 on each cancer individually. The correlation between paired tumor and matched normal samples from the same patient was modeled using the duplicateCorrelation() function, allowing for appropriate handling of intra-patient dependencies. Differential testing was performed using limma’s empirical Bayes (EB) moderation, which stabilizes variance estimates and improves statistical power, particularly in datasets with limited sample sizes. To further increase robustness, the EB procedure was carried out with robust = TRUE to reduce the influence of potential outliers and trend = TRUE to account for mean–variance relationships in the metabolic activity data. For the Zhou2020 and Hu2025 proteomics data, standard limma differential analysis was performed whereever there was not enough paired data. In the Zhou2020 and Hu2025 datasets, CellFie task metabolic activity scores were log2 transformed before limma analysis, and only metabolic tasks with at least two quantified scores in each sample group (normal and tumor) were included in limma analysis. For the twelve cancer types in the Hu2025 data, we also controlled for batch effects when multiple batches were present within a single cancer type. Benjamini-Hochberg correction was applied to control the false discovery rate.

### Differential analysis of metabolic-related protein abundance between cancer and normal samples

CellFie was originally designed for bulk RNA-seq with extensive gene coverage; thus, some metabolic tasks in proteomics data may have insufficient protein measurements for stable activity scores. To validate CellFie-derived patterns, we performed paired differential expression analysis for each metabolic class at the protein level using the same limma-based approach. This analysis included only paired samples and proteins quantified in at least two samples per group (normal and tumor).

### Gene co-expression network analysis

Gene co-expression network analysis was performed to uncover coordinated regulation and identify key hub genes driving metabolic rewiring in cancer. For each KEGG metabolic class, pairwise Spearman correlations among member genes were computed across TCGA samples within each cancer type and averaged to generate a co-expression matrix. Gene connectivity was calculated as the sum of absolute mean correlations excluding self-correlations, with the top 1% most connected genes (minimum three per class) defined as hubs. Edges with absolute mean correlation > 0.6 were retained to construct undirected weighted networks using igraph v2.0.3.

### Multi-omics integration

To integrate the two proteomics datasets with the TCGA cohort, we employed the iNMF method from rliger v1.0.0 [48] for a joint analysis of KEGG-derived metabolic activity scores. The analysis was restricted to pathways associated with glycan and nucleotide metabolism.

All analyses were conducted in R v4.4.3. We developed the **parseMetab** R package (v1.0.4, (https://github.com/Maj18/parseMetab, https://doi.org/10.5281/zenodo.19153922) to streamline the workflow, enabling inference and analysis of metabolic activity from omics data, differential and network analyses, and cohort-level visualizations. All computations were performed within a Docker container (Docker Hub ID: yuanli202004/cancer:v2.0.2) to ensure reproducibility and consistent computational environments. All R code required to reproduce the analyses is available at https://github.com/Maj18/cancer.

## Acknowledgements

We thank Lund university library for supporting open access publications. The authors thank Carl Brunius from NBIS for valuable comments and suggestions on an earlier draft of this manuscript.

## Ethics approval and consent to participate

This study involved secondary analysis of publicly available, de-identified proteomic, transcriptomics and spatial transcriptomics datasets. No new human participants or animal subjects were recruited, and therefore ethics approval and consent to participate were not required.

## Consent for publication

Not applicable.

## Availability of data and materials

The dataset(s) supporting the conclusions of this article is(are) included within the article (and its additional file(s)).

## Authors’ contributions

Y.L. conceived and designed the study; Y.L., Y.B. and Q.C. analyzed the data, prepared figures and/or tables. Y.L., Y.B. and Q.C. wrote the paper. All authors read and approved the final manuscript.

## Additional files

**Fig.S1.pdf: Fig. S1. Summary of metabolic activity and gene-level characteristics across pan-cancer datasets.** (A-D, K-L) Boxplots showing mean metabolic activity per task for each cancer type, with one panel per dataset. (E-H, M-N) Boxplots showing mean metabolic activity per task for each metabolic class, with one panel per dataset. (I-J) Violin plots of mean protein expression for each metabolic class for the two proteomics datasets. (O-S) Boxplots of task/pathway size (number of genes per task) across metabolic classes for different datasets. (T-U) Violin plots showing the distribution of moderated *t* statistics from pathway-level differential analyses for each metabolic class in Hu2025_KEGG and Zhou2020_KEGG. (V-W) Circular barplots showing the number of metabolic genes for each cancer type within each dataset.

**Fig.S2.pdf: Fig. S2. Pan-cancer metabolic dysregulation across independent pan-cancer datasets from two distinct omics modalities.** Dot plots of significant task-level dysregulation across cancer types in the Zhou2020 (A, adj. P < 0.1) and SMTdb (B, P.value < 0.1) data. Color represents moderated *t* statistics, and dot size reflects (adjusted) *p* values.

**Fig.S3.pdf: Fig. S3. Pan-cancer metabolic dysregulation across independent pan-cancer datasets from two KEGG-based proteomics data.** Dot plots of significant pathway-level dysregulation across cancer types in the Hu2025_KEGG (A, adj. P < 0.1) and Zhou_KEGG (B, P. adj. P < 0.05) data. Color represents moderated *t* statistics, and dot size reflects (adjusted) *p* values.

**Fig.S4.pdf: Fig. S4. Gene co-expression networks for different metabolic classes.** Networks nodes represent genes, sized by normalized connectivity, with the top 1% most connected genes (≥3 per class) highlighted as hubs (red). Edges show absolute correlation > 0.6, width scaled by correlation strength; grey edges indicate positive correlations and blue edges indicate negative correlations. Networks were visualized using a Fruchterman–Reingold layout

**TableS1.xlsx: Table S1. Summary and sample information for pan-cancer datasets analyzed in this study.** The summary sheet provides an overview of all included datasets and their corresponding omics modalities, cancer types, sample counts and identified task/pathway/protein/gene numbers. Additional sheets contain detailed sample-level information for each dataset, including sample identifiers, tissue type (tumor or normal), and associated clinical annotations.

**TableS2.xlsx: Table S2. Metabolic activities inferred from proteomics, bulk transcriptomics, and spatial transcriptomics across pan-cancer datasets.**

**TableS3.xlsx: Table S3. Pan-cancer metabolic dysregulation results.** This Excel file contains multiple sheets showing differential analyses across cancer types and datasets. Task/Pathway-level sheets (MA_*): Results for metabolic tasks or pathways for each dataset and cancer type. Protein-level sheets (Protein_*): Protein-level differential analysis within metabolic classes for each dataset. Sheet names indicate the level (task/pathway or protein), dataset, and cancer type/class for easy reference.

